# Resistance Gene-Directed Genome Mining of 50 Aspergillus species

**DOI:** 10.1101/457903

**Authors:** Inge Kjærbølling, Tammi Vesth, Mikael R. Andersen

**Affiliations:** Technical University of Denmark, Department of Biotechnology and Biomedicine, Kongens Lyngby, Denmark

**Author notes:** Address correspondence to Mikael R. Andersen.

**Keywords:** secondary metabolism, resistance, genome mining, Aspergillus

## Abstract

Fungal secondary metabolites are a rich source of valuable natural products. Genome sequencing have revealed an enormous potential from predicted biosynthetic gene clusters. It is however currently a time consuming task and an unfeasible task to characterize all biosynthetic gene cluster and to identify possible uses of the compounds. A rational approach is needed to identify promising gene clusters responsible for producing valuable compounds. Several valuable bioactive clusters have been shown to include a resistance gene which is a paralog of the target gene inhibited by the compound. This mechanism can be used to design a rational approach selecting those clusters.

We have developed a pipeline FRIGG (Fungal Resistance Gene-directed Genome mining) identifying putative resistance genes found in biosynthetic gene clusters based on homology patterns of the cluster genes. The FRIGG pipeline has been run using 51 *Aspergillus* and *Penicillium* genomes, identifying 72 unique protein families with putative resistance genes using various settings in the pipeline. The pipeline was also able to identify the characterized resistance gene *inpE* from the Fellutamide B cluster thereby validating the approach.

We have successfully developed an approach identifying putative valuable bio-active clusters based on a specific resistance mechanism. This approach will be highly useful as an ever increasing amount of genomic data becomes available — the art of identifying and selecting clusters producing novel valuable compounds will only become more crucial.

**Importance:** Species belonging to the *Aspergillus* genus are known to produce a large number of secondary metabolites, some of these compounds are bioactive and used as pharmaceuticals such as penicillin, cyclosporin and statin. With whole genome sequencing it became apparent that the genetic potential for secondary metabolite production is much bigger than expected. As an increasing number of species are whole genome sequenced an immense number of secondary metabolite genes are predicted and the question of how to selectively identify novel bioactive compounds from this information arises. To address this question, we have created a pipeline identifying genes likely involved in the production of bioactive compounds based on a resistance gene hypothesis approach.

## Introduction

Fungal secondary metabolites are a rich source of bio-active compounds including important pharmaceuticals such as penicillin, cyclosporin and statin (1). When the first fungal genomes were sequenced, it became clear that the genomes harbour a higher number of secondary metabolite gene clusters than the number of characterized secondary metabolites thus revealing a much larger potential (2, 3, 4, 1). The number of sequenced genomes is ever increasing mainly due to large sequencing efforts such as the 1000 Fungal Genomes Project of the Department of Energy Joint Genomes Initiative (http://1000.fungalgenomes.org/home/) and the 300 Aspergillus genome project (5, 6) and therefore the number of predicted secondary metabolite gene clusters is steadily increasing.

Despite progress in molecular tools and methods for characterization of secondary metabolite gene clusters, it is still a time-consuming task, making it unfeasible to investigate all predicted secondary metabolite gene clusters. Therefore only a small fraction of the predicted clusters are characterized and investigated experimentally. With the plethora of predicted secondary metabolite gene clusters (clusters) and the aim of discovering novel bio-active compounds useful as drugs, the question emerges: How do we select the most interesting predicted clusters producing potential valuable drugs such as anti-fungicides, anti-cancer drugs and anti-microbial compounds? To meet this need we have created a pipeline FRIGG (Fungal Resistance Gene-directed Genome mining) identifying clusters producing likely bio-active compounds based on resistance genes. Many bio-active compounds are toxic compounds also impairing the organism that synthesize them by inhibiting essential functions, therefore a self-resistance mechanism is needed in order to survive (7, 8, 9). One known self-resistance mechanism is the duplication of the target gene, where the second version is resistant towards the compound and this second resistant version is most often found as part of the biosynthetic gene cluster producing the toxic compound. This mechanism has been seen in several bacterial instances such as novobiocin (10) and pentalenolactone (11, 12).

More recently this resistance mechanism has also been identified in fungi. Mycophenolic acid (MPA) is produced by *Penicillium brevicompactum* and it inhibits inosine-5′-monophosphate dehydrogenase (IMPDH) which is the rate limiting step in guanine synthesis. The biosynthetic cluster of MPA revealed an additional copy of IMPDH which is insensitive to MPA thus conferring resistance, Figure 1A (13, 14, 15). Another example is Fellutamide B produced by *A. nidulans* which is a proteasome inhibitor. Within the biosynthetic gene cluster a gene, *inpE*, encoding a proteasome subunit is located and it was shown that this gene confers resistance, Figure 1B (16).

**FIG 1.**
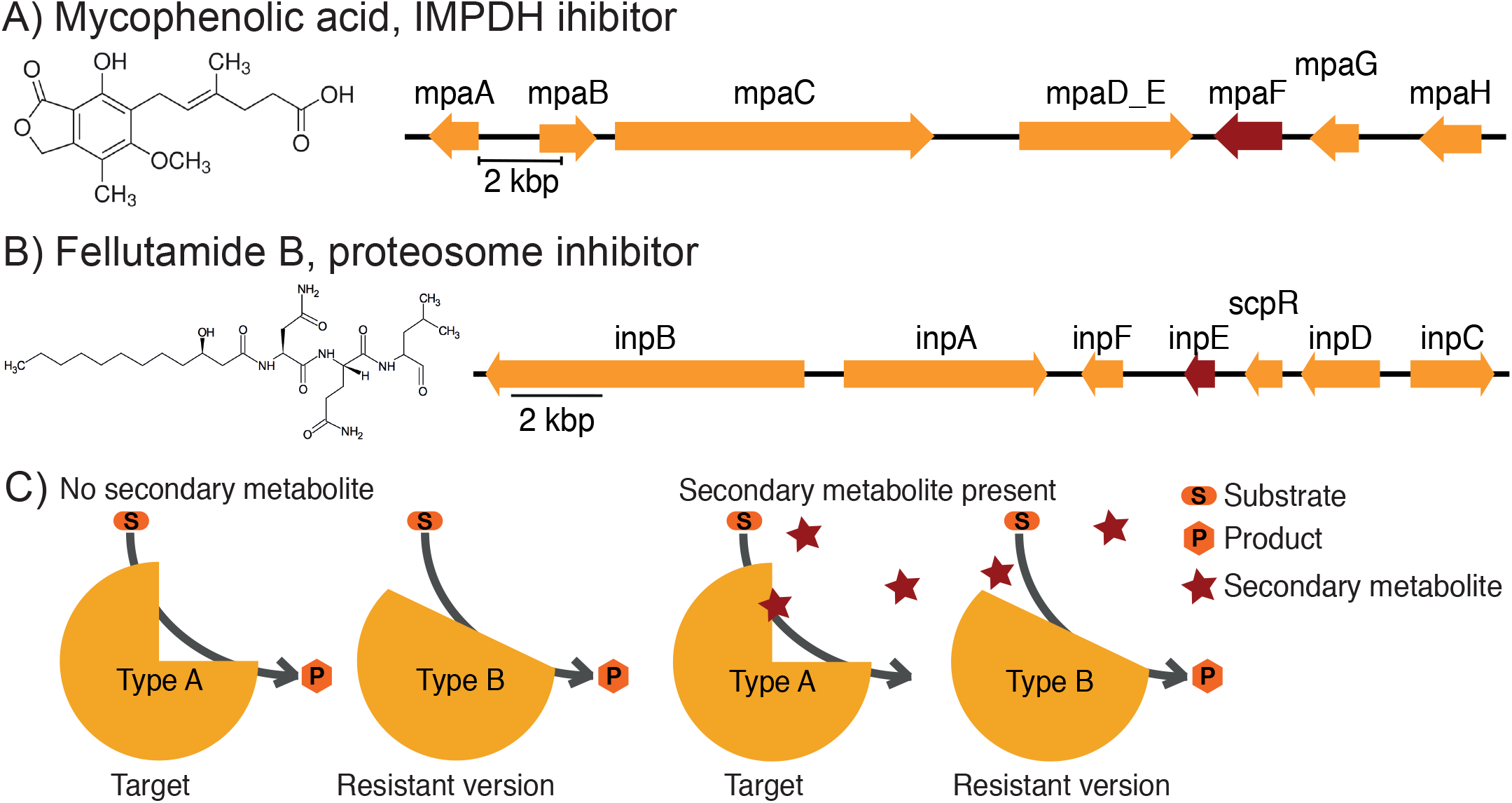
Resistance mechanism. A) Mycophenolic acid chemical structure and biosynthetic cluster with known resistance gene mpaF (highligted in red) which is a Inosine-5′-monophosphate dehydrogenase (IMPDH) inhibitor. B) Chemical struture of Fellutamide B and overview of the biosynthetic cluster including the resistance gene inpE (highlighted in red) which is a proteasome inhibitor. C) Illustration of the resistance mechanism used by some toxin producers. The secondary metabolite is a toxin which inhibits an essential enzyme - the target of the compound. Within the cluster responsible for producing the toxin a copy of the target gene is found, this version is still functioning despite the compounds presence and hence makes the organism self-resistant.

An illustration of the general mechanism can be seen on Figure 1C, where two versions of an enzyme is present and one version is affected by the secondary metabolite (the target) whereas the other version is slightly different, but still with the same function, is not inhibited by the secondary metabolite (the resistance gene). Even though only a few examples of this mechanism have been identified and verified in filamentous fungi so far (13, 16, 17, 18), it is possible that this resistance mechanism is more widely distributed. We have therefore developed a Fungal Resistance Gene-directed Genome mining (FRIGG) pipeline identifying putative bio-active clusters with resistance genes. The aim of the pipeline is to identify bio-active clusters in a targeted manner thus providing a way of selecting the most interesting predicted clusters producing potential valuable drugs from whole genome sequences.

The immediate advantage of the FRIGG pipeline is that highly likely bioactive clusters are identified. Another major advantage of using this approach is that the target of the compound is inherently known and hence the mode of action. Knowing the target saves a lot of time since linking the compound to the target is extreme difficult and time consuming and several regular drug discovery steps can be eliminated since possible uses of the compound are known from the beginning.

## Results

### Pipeline set-up and Input data

We were interested in creating a pipeline identifying secondary metabolite gene clusters (clusters) containing possible resistance genes from whole genome sequencing data. In order to do this we based the pipeline on the assumption that the resistance gene is found within a cluster and is a copy – a paralog – of an essential gene. This pattern and the resulting resistance mechanism has previously been described in two different cases in fungi (13, 16).

We have used complete and draft quality whole genome sequences, mainly from the 300 *Aspergillus* sequencing project (5, 6). The input for the pipeline was chosen to consists of three different types of data derived from the whole genome sequence data: 1) predicted genes/proteins and functionally annotated proteins, for the functional annotation we used InterPro (19, 20). 2) Predictions of secondary metabolite gene clusters, for this purpose we used a re-implementation of SMURF (21) as described in Vesth et al. (6). 3) Groups of homologous protein sequences. We used a pipeline designed for *Aspergillus* data creating homologous protein families based on single linkage of bidirectional BLASTp hits (as described in (6)). It is assumed that proteins with similar, although not necessarily identical, function will be clustered into the same protein family. For our purpose this is useful, as resistance and target genes will be grouped into one family.

Using the described input the pipeline consists of a number of filtration steps designed to identify the most likely candidate clusters containing potential resistance genes, Figure Figure 2. Several steps in the pipeline are designed to deal with and/or minimize the effect of possible errors in assembly and annotation due either to inherent errors in sequencing technologies or errors caused by the assembly and sequence quality of draft genomes. Several options have been added to the pipeline to allow the user to set the filters to allow for more or less noise caused by errors. Besides mitigating errors the filtering steps are also implemented to deal with biological diversity and differences. Here we present the results of the pipeline using an extensive test dataset of 51 *Penicillium* and *Aspergillus* species containing a total of 3,276 predicted clusters and 26,551 secondary metabolite genes. The goal of the pipeline is to identify clusters containing resistance genes, assuming that the resistance genes are copies of essential genes. In this context, an essential gene is defined as a gene that has homologs in all the species included in the analysis.

**FIG 2.**
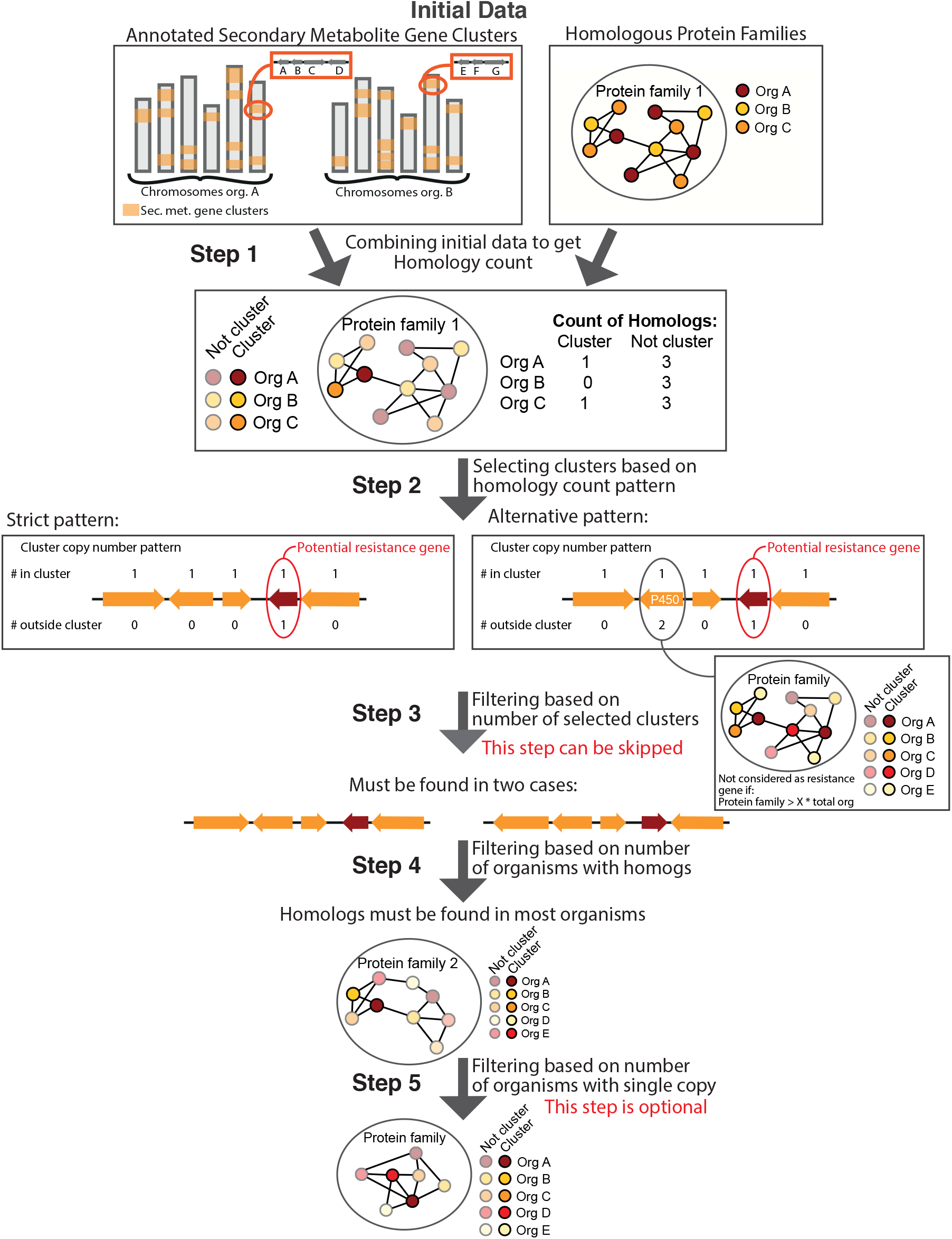
Overview of the pipeline. Overview of the pipeline illustrating the initial data of predicted secondary metabolite gene clusters and homologous protein families. Step 1 – The initial data is combined to generate counts of how many homologs are found in each organisms andhow many are within predicted clusters. Step 2 – Clusters are selected based on specific homology count patterns, either ‘strict’ where only one gene can have a homolog outside the cluster or ‘Alternative’ where genes belonging to large protein families are allowed. Step 3 – Filtering of the selected clusters based on other clusters having a homologous resistance gene, this step is optional. Step 4 – Filtering based on the majority of the organisms should have a homolog of the resistance/target genes. Step 5 – Filtering based on the number of organism that should only have a single homolog.

### Homolog count and ‘Strict’ cluster selection

The first step in the pipeline is to couple the homologous protein families to the predicted cluster genes, Figure Figure 2 step 1. Following, the number of homologs – homology count – for each secondary metabolite gene in each organism/genome is identified. In addition, the number of homologs found in predicted clusters are recorded.

Next, clusters with potential resistance genes are selected based on a specific pattern of homology counts, Figure Figure 2 step 2. In this step the user can select various levels of stringency for the selection pattern.

The most strict and simple selection pattern (Figure Figure 2 step 2 left) identifies clusters where only one of the genes have a homolog in the genome, it can have only one such homolog, and this homolog must not be part of another cluster. The gene with the homolog is the presumed resistance gene and the homolog outside the cluster is the presumed target gene. Using this selection criteria on our test dataset, 262 clusters are identified, divided into 141 potential resistance genes protein families, Figure Figure 3A (dark turquoise). This corresponds to 8% of the total clusters in the data set.

**FIG 3.**
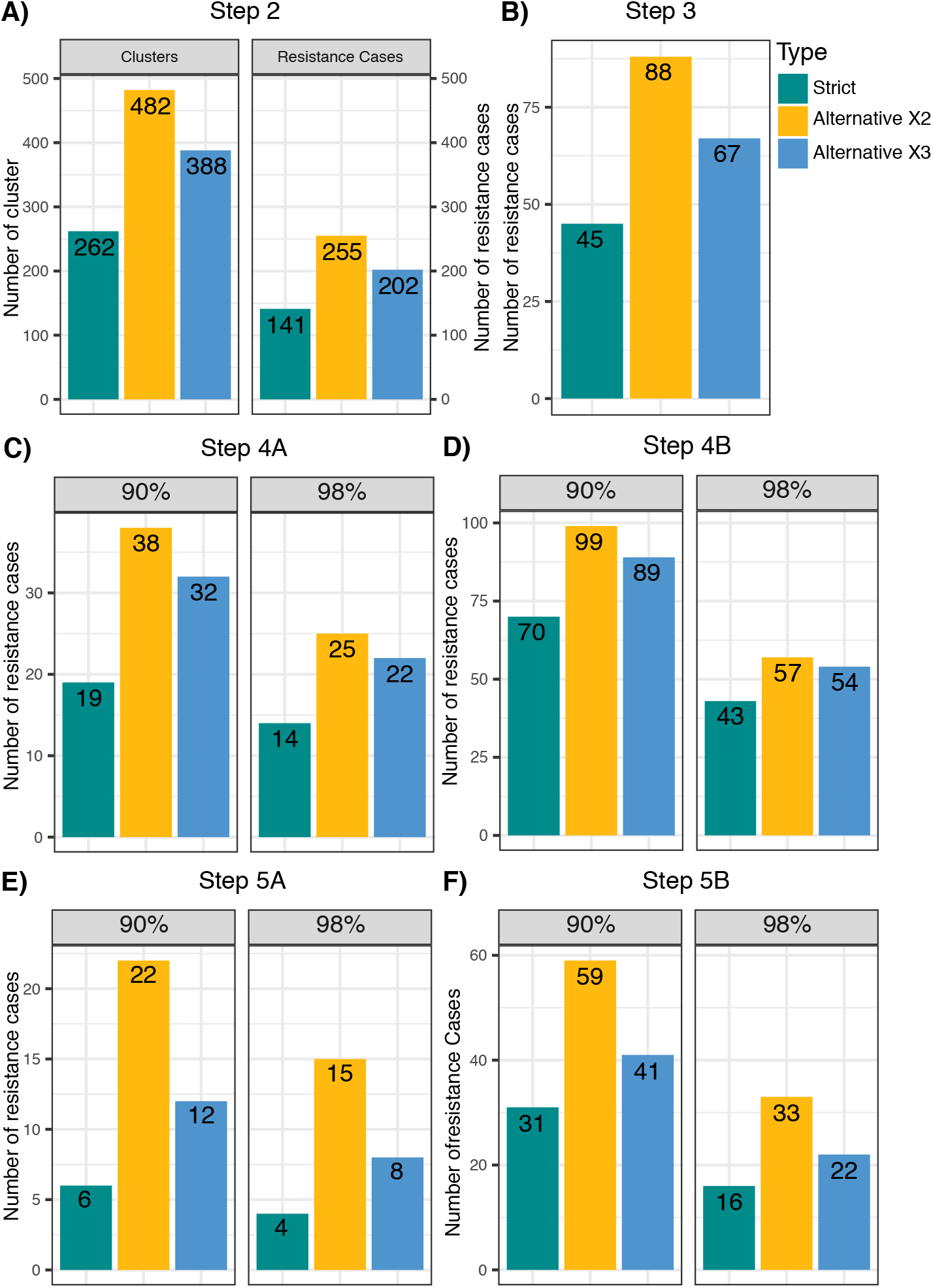
Number of cases after each filtering step using various settings. A) the number of cluster selected for each of the selections patterns and the number of resistance cases. B) The number of resistance cases after step 3 ‐ the resistance protein family has to be found in two selected clusters. C) The number of resistance cases after step 4 ‐ the resistance protein family have to have homologs in 90 or 98% of the organisms in the dataset. D) The number of resistance cases after step 4 but having skipped step 3, again both for 90 and 98% of the organisms. E) The number of resistance cases after step 5 ‐ the resistance protein family have to be found as single copy in 50% of the species. F) The number of resistance cases after step 5 but having skipped step 3.

This ‘strict’ selection pattern is very restrictive: if any one of the other cluster genes have a homolog anywhere in the genome, the cluster will be filtered away. Many clusters contain tailoring enzymes with common functions such as P450 or methyltransferases, these can be thought of as “household” functions in secondary metabolism. Clusters with common tailoring functions will therefore frequently have several homologs resulting in high homology counts and the cluster will therefore fall out of the ‘strict’ selection pattern. Another effect of the filtering is that the selected clusters often contain fewer genes (average size 6.4 genes vs. 8.2 for the total data set). The bigger the cluster, the more likely it is that there are several genes with homologs in the genome. It is likely that clusters containing potential resistance genes are missed due to the strict selection criteria. Conversely, the share of bioactive clusters is increased after the selection.

### ‘Alternative’ cluster selection pattern

To increase the number of selected clusters with potential resistance genes, we wanted to create an alternative selection pattern allowing the presence of more tailoring enzymes and larger cluster sizes. In order to generate reasonable alternative selection criteria, the cluster genes in the dataset were investigated. The most common InterPro domains (annotated in at least 1000 cluster genes) were selected, the protein families belonging to each InterPro domain was identified and the size of the protein families were noted. In the additional file 5 a violin plot illustrates the selected InterPro domains on the x-axis and the sizes of the protein families with at least one protein with this domain are on the y-axis. There are many protein families belonging to each InterPro domain ranging in size from 1 to 475 proteins.

Protein families with more than 100 members will most likely have several homologs in each organism and every time they appear in a cluster, the cluster will be discarded using the ‘strict’ selection pattern. To avoid this, an ‘alternative’ selection pattern was created disregarding large protein families as potential resistance genes and instead allowing the proteins belonging to large protein families to have homologs in the organism, Figure Figure 2 step 2 ‘alternative’ pattern.

The size of the protein families will change depending on the data (e.g. the number of genomes included and how closely related the species are). The selection was therefore designed to be dependent on the number of organisms in the data set. In order to determine this, we defined a metric: If the protein family is larger than the total number of organisms in the dataset multiplied by X, where X is a user input, then the gene is not considered as a potential resistance gene. Instead it is allowed to have homologs and the cluster can still be selected, if another gene meets the requirements for a resistance gene and the rest of the genes either have no homologs or belong to a large protein family. The alternative pattern under step 2 in Figure Figure 2 illustrates the copy number pattern.

In the less strict selection method, the potential resistance gene is thus still only allowed to have one homolog outside the cluster and there is only one gene that can have this pattern, however the other genes in the cluster are allowed to have more homologs if their protein family is larger than: *X_Input_* × *Number of organisms*. If X is set to 2 then the cutoff in the illustration (step 2 in Figure Figure 2) would be 2*5 organisms and the protein family illustrated has 11 members and hence would be allowed in the pattern. If 3 is selected instead the cutoff would be 12 and the illustrated protein family is not big enough and this cluster would fall out.

We have used two and three as input thereby disregarding protein families with more than 102 or 153 members respectively. With this selection criteria 482 and 388 clusters are identified respectively which corresponds to an 84% and 48% increase compared to the initial ‘strict’ selection criteria, Figure Figure 3A. Of these clusters there are 255 and 202 different potential resistance gene families, corresponding to an increase of 81% and 43% respectively, showing that the ‘strict’ measurement is indeed sensitive to large protein families.

### Filtering

As mentioned above, genome data contains multiple types of errors such as incomplete genomes, gene calling and secondary metabolite gene cluster prediction errors. Incomplete genomes and gene calling errors are likely to casue false negatives in our pipeline while cluster prediction errors can cause both false positives and negatives depending on if the prediction algorithm over or under predicts the size of the cluster. Even with the current rate of technological advancement, data errors will most likely be a problem for some time to come and therefore we have created filtering steps to deal with these shortcomings in the data. Another reason for filtering is to identify the most likely resistance genes and hence filtering steps have also been implemented to decrease false positives.

### Step 3 – Filtering the number of clusters

A filtering step is implemented in order to mitigate secondary metabolite prediction errors (Figure Figure 2 step 3). With the assumption that if two of the selected clusters have potential resistance genes belonging to the same protein family, then it is less likely that it is an error and more likely that the resistance gene belongs to the cluster. The filtering step selects only the clusters where at least one other identified cluster has a potential resistance gene belonging to the same protein family. As such, the members of the potential resistance gene protein family have to be found in at least two selected clusters.

Using these filtering criteria, 45, 88, and 67 of the previous identified potential resistance genes protein families are left for the ‘strict’, ‘alternative’ *X_input_* 2 and 3 which correspond to 32-35% of the initial selected potential resistance gene cases, Figure Figure 3B. As such about a third of the initially identified clusters share resistance gene with another identified cluster and these are therefore more likely not to be prediction errors.

### Step 4 – Filtering the number of organisms with homologs

The assumption behind this step is that the resistance genes should preferentially belong to a protein family with an essential function, and thus it should have protein members in all the species. The filtering step has two purposes, first it makes it more likely that it is a resistance gene if there are homologs in all species and second it will be a more widely useful bioactive compound if the target is conserved in many species.

This filtering step removes clusters where the putative resistance gene have homologs in less than a certain percentage of the organisms, Figure Figure 2 step 4. The user can select percentages from 100 to 90 to select the most likely resistance genes and best targets. The possibility of choosing lower than 100% was implemented to allow for some data and prediction errors such as incomplete genomes and imperfect gene annotations. Another reason for choosing lower than 100% is to allow some species to have a different mechanism and thus not the target gene. Here we show the effect of selecting 98% and 90% of the organism on the number of cases selected. If selecting the most restrictive setting where the homologs of the resistance gene has to be found in 98% of the organism, then there are 14, 25 and 22 potential resistance gene cases of the ‘strict’, ‘alternative’ *X_input_* 2 and 3 respectively, Figure Figure 3C. While setting the percentage of organisms to 90% there are 19, 38 and 32 potential resistance gene cases, Figure Figure 3C.

If one chooses not to use step 3 (based on at least two selected clusters having the potential resistance gene), but only employ step 4 (based on the number of organisms homologs of the potential resistance gene should be found in) more clusters are selected, see Figure Figure 3D. Using the setting of potential resistance genes having homologs in 98% of the organisms 43, 57 and 54 cases are identified based on the homology count pattern of ‘strict’, ‘alternative’ *X_input_* 2 and 3, respectively. While employing only 90% of the organisms should have homologs of the potential resistance genes, 70, 99 and 89 are cases were detected for ‘strict’, ‘alternative’ *X_input_* 2 and 3.

### Step 5 – Filtering the number of organisms with single copies

The final filtering step is again related to the organisms in which the putative resistance gene have homologs. The assumption here is that the target gene should be one essential gene and therefore the majority of the species should only have one copy.

This filtering step therefore removes protein families where more than 50% of the organisms have more than one homolog of the putative resistance gene, Figure Figure 2 step 5. This step is optional and can be employed or not as the user sees fit. The effect of adding this selection criteria can bee seen in Figure Figure 3E and F.

### Pipeline output

The primary output of this pipeline is a list of protein families with potential resistance genes and fasta files containing all the proteins belonging to the identified family. The header of each entry includes the name of the organism, the section it belongs to, the protein id, the number of copies found in the organism and if it is in a selected cluster (StrictClust), a predicted cluster (Clust) or somewhere else in the genome (0) which is followed by the amino acid sequence. The cases that come out of the pipeline depends on the different settings and each identified case is independent of the others. If a high experimental capacity is available one option is to test all these cases, if the experimental capacity is limited further analysis is needed to evaluate and select which cases to work with.

In this example with 51 species, we get 12 different outputs after step 5 using each of the various settings. Narrowing down from 3,276 clusters we have identified 72 unique putative resistance gene families using all the various settings (Table 5). Of these 4 potential resistance gene families are found every time independent on which setting, while 19 are found only once using a specific set of variables.

With each filtering step, the number of clusters and putative resistance genes decreases, but the share of likely bioactive clusters and true resistance genes increases.

### Identifying potential bioactive clusters

Even with the described selection criteria, the pipeline still produces more candidate clusters than it would be feasible for most academic labs to verify experimentally. Once the pipeline has been run and potential resistance gene cases have been identified, further analysis is needed to select the most promising candidates for experimental verification.

In order to do this efficiently, we use a combination of principal component analysis (PCA), phylogenetic analysis, functional annotation, and comparison to the NCBI database using BLASTp in order to gain more knowledge about the potential cases (examples of the PCA and phylogenetic trees can be seen in Figure Figure 4 and additional Figure 5 and 5).

The functional annotation and BLAST analysis are included to add to the understanding of the potential resistance genes; to examine if it has a known function, if the function is essential, and if it would be a good drug target.

**FIG 4.**
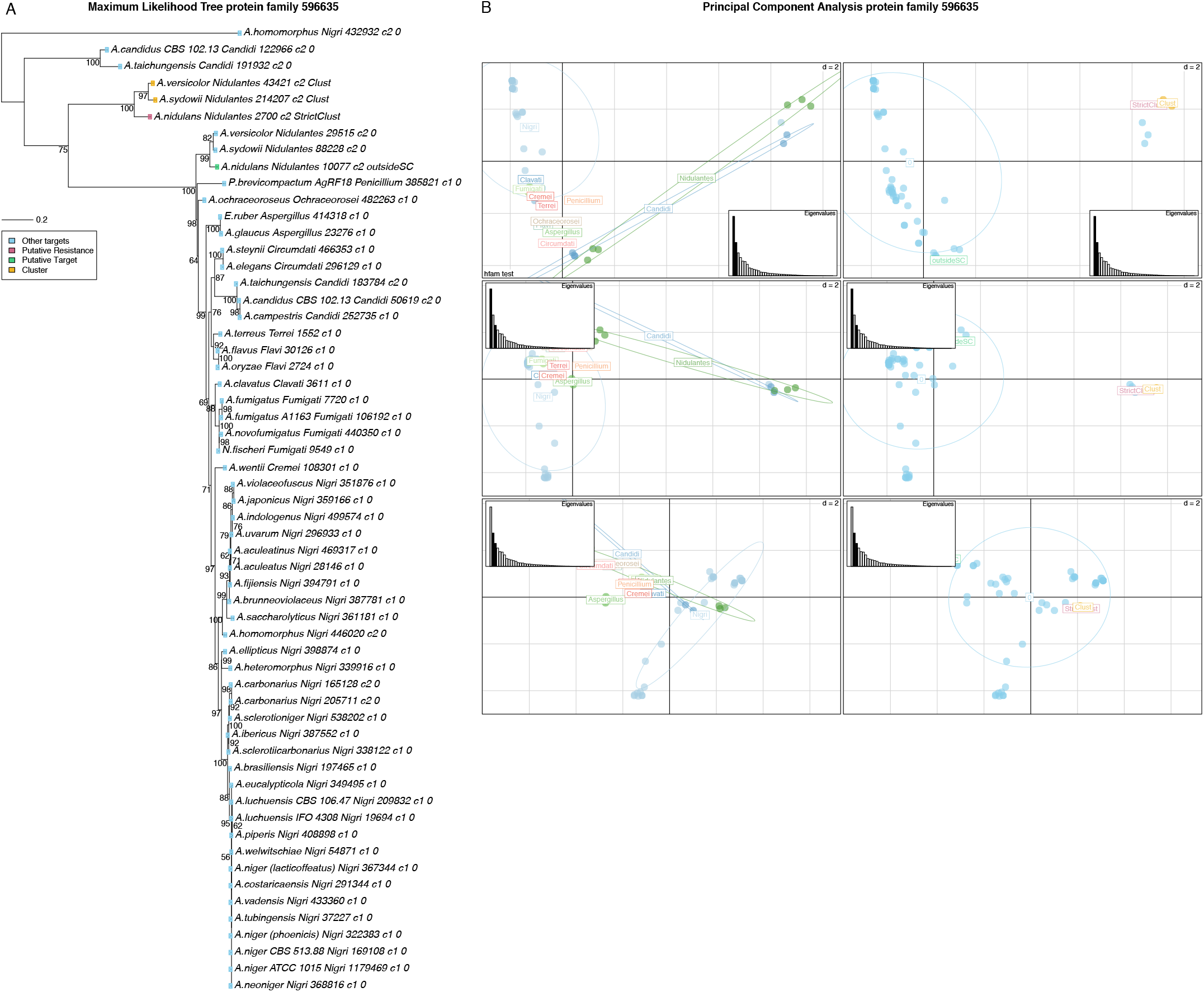
Phylogenetic tree and principal component analysis of protein family 596635 including the Fellutamide B resistance gene. A) Phylogenetic tree of the the protein family (596635) containing the Fellutamide B resistance gene, (protein id 2700) from *A. nidulans*. The phylogenetic tree is a maximum likelihood tree and was created with 500 bootstraps. The identified resistance gene is highlighted with red, the corresponding target is in green, other cluster genes are in yellow and gene not found in clusters are in blue. B) Principal component analysis of the protein family 596635 containing the Fellutamide B resistance gene, (protein id 2700) from *A. nidulans*. The panels to the left is colored based on the sections the proteins belong to while the panels to the right is colored based on if the protein if found in a a selected cluster (StrictClust-red), a predicted cluster (Clust-yellow), not in a cluster (0-blue) and the homolg to the ones found in selected clusters (outsideSC-green).

The PCA and phylogenetic trees are used to illustrate the evolutionary relationship between the proteins in the protein family. Resistance genes are expected to be a duplication of a target gene which is essential. Essential genes are under similar evolutionary pressure in all the species and are therefore closely related. The resistance gene however, is under a different pressure, potentially expressed under a different subset of conditions, which is expected to be reflected in the sequence and hence the analysis where resistance genes should form a separate group compared to the target genes. To perform these two analysis the protein sequences from the protein families were aligned using clustalo (22) and trimmed using Gblocks (23, 24).

As mentioned in the introduction, there are some known clusters containing a verified resistance gene, this includes the mycophenolic acid cluster in *P. brevicompactum* and the fellutamide B cluster from *A. nidulans*. Both clusters were included in this study as validation controls to check if the known resistance genes would be identified using our pipeline.

The Fellutamide B cluster is identified in all the outputs where step 3 was skipped, since this was the only cluster with this resistance gene and the selected homology count pattern. Thus when filtering for more reliable cluster predictions it falls out of the analysis.

The potential resistance and target genes are exposed to different evolutionary pressure. This is expected to be reflected in the evolutionary pattern where the essential target genes (under high pressure) will be closely related whereas the resistance genes will show more variation. To investigate if this is the case a phylogenetic tree and a principal component (PCA) analysis was performed for the protein family containing the fellutamide B resistance gene (596635), Figure 4. The PCA shows a clear distinction between the resistance genes found in clusters and the other essential target genes, Figure 4B. The phylogenetic tree (Figure 4A) shows two clear groupings: one big closely related group with all the target genes (having 0 at the end of the label), and one small containing the resistance gene. The target genes are ordered in their phylogenetic groups, so species from the same section cluster together. The resistance gene from *A. nidulans* is clustered with a protein from *A. sydowii* and *A. versicolor*, which both are found in clusters, indicating that these most likely also function as resistance genes and potentially can produce Fellutamide B or a derivative or a different compound attacking the same target. *A. versicolor* is known to produce fellutamide C and F (25, 26), whereas *A. sydowii* to our knowledge has not been reported to produce derivatives of fellutamide. The clusters in *A. versicolor* and *A. sydowii* are similar to the *A. nidulans* fellutamide B cluster with similar backbone enzymes (≥ 70%) and tailoring enzymes, but the *A. versicolor* and *A. sydowii* predicted clusters are bigger consisting of 19 genes, 7 more than the *A. nidulans* cluster. Three of the extra genes have homologs in the genome and belong to small protein families which is why these clusters are not identified in the first steps of the pipeline.

Near the resistance genes in the phylogenetic tree, there are also three other genes found in *A. homomorphus*, *A. tachungensis* and *A. candidus* which are extra copies of the target genes, but they are not predicted to be in clusters. One explanation for the extra copies not found in clusters could be cluster prediction errors. We therefore investigated if *A. homomorphus*, *A. tachungensis* and *A. candidus* had a cluster similar to the *A. nidulans* fellutamide B cluster, but no BLASTp hits of the backbone genes had identity above 37% indicating that they most likely cannot produce any derivatives of fellutamide. Another explanation for the extra copy could be that it functions as a defence mechanism protecting against other species producing fellutamide, which could be useful if the species naturally grow near fellutamide producing species.

The cluster of mycophenolic acid did not turn up in our outputs, so we investigated the cluster further to understand why. The predicted cluster containing the PKS responsible for producing the core compound of mycophenolic acid consists of four genes; The PKS, a P450, a methyltransferase and the resistance gene. The PKS and p450 are only found in one copy and the resistance gene has one identified homolog as expected. The methyltransferase however also has a homolog in the genome. As the size of this protein family is only 23 members, it is not disregarded in the alternative pattern and the cluster is therefore filtered away in step 2. This shows that we do loose some good cases along the steps in the pipeline which illustrates the importance of running the pipeline with multiple settings and inspecting the output carefully. The pipeline is highly sensitive to the number of organisms included and the settings should be used keeping this limitation in mind.

### Novel putative resistance gene

In addition to the known clusters with resistance genes, several uncharacterized clusters were also identified containing putative resistance genes. We will here focus on one example where the potential resistance gene is found in clusters in *A. oryzae* and *A. flavus* (protein family 597268). The PCA analysis showed a very clear picture of the resistance genes falling outside the group of target genes, the same is seen in the phylogenetic tree, where the target genes also follow the expected phylogeny with species from the same section grouping together, Additional Figure 5 and 5. Besides the identified putative resistance genes in *A. oryzae* and *A. flavus* several other species (*A. wentii*, *A. piperis*, *A. candidus*, *A. taichungensis*, *A. campestris*, *A. novofumigatus* and *P. brevicompactum*) have an additional gene but these are not found in predicted clusters. The backbone genes from *A. oryzae* and *A. flavus* have no hits in those species which could mean that the species carries the resistance gene but do not produce the compound.

The predicted clusters in *A. oryzae* and *A. flavus* both consists of four genes: an acetyltransferase (IPR000182, IPR016181), the predicted reistance gene, a NRPS-like synthetase and a gene belonging to the major facilitator superfamily (IPR011701, IPR007114). The putative resistance gene has annotations involved in ‘Signal transduction response regulator, receiver region’ and ‘Signal transduction histidine kinase, core’ (IPR001789, IPR005467). Using BLASTp to investigate the function of the putative resistance gene, the top hits have functions like: ‘two-component osmosensing histidine kinase (Bos1)’ (RAQ55620.1). Based on this information it seem likely we have identified a cluster containing a previously unidentified resistance gene where the compound inhibits a histidine kinase. In order to fully determine this, this will have to be experimentally validated in the future.

## Discussion

The pipeline was designed to identify secondary metabolite gene clusters (clusters) containing potential resistance genes. It is a delicate balance of filtering away as many clusters as possible to narrow down the field to the best candidates, while keeping as many clusters with potential resistance gene as possible.

We have chosen an approach, where we make no assumptions about the function or makeup of resistance genes besides being homologs of a gene shared by most organisms in the data set. Another approach could be to screen for specific classes of essential genes or resistance targets within predicted clusters. This approach was used in another study where 86 bacterial *Salinispora* genomes were mined for duplicated genes involved in central metabolism co-localizing with clusters. Clusters containing putative fatty acid synthase resistance genes were identified and these were shown to be involved in the biosynthesis of thiotetronic acid natural products, including thiolactomycin which is a well-known fatty acid synthase inhibitor (27). This approach builds on knowledge of house-keeping genes thus requiring extensive knowledge of the primary metabolism. In filamentous fungi there is still a lot of primary processes which are not characterized, therefore there is a risk of missing interesting resistance genes using that approach. To avoid this, we chose to use a wider approach based only on the homology copy number pattern of predicted secondary metabolism genes and not based on the functions. The underlying assumption is that our pipeline identifies conserved household genes with a homolog in a cluster. Using our approach, we avoid limiting the search space to only known mechanisms thereby making it possible to find new essential mechanisms and drug targets. Our non-functional-impelled approach can be used both on organisms with little knowledge and extensive knowledge of the primary metabolism. In well characterized species our approach is also useful since it is likely both to identify known household genes but also new uncharacterized household genes. Finally, the setup makes it possible to search the identified clusters afterwards with a criterion such as the presence of primary metabolism genes. The pipeline has been designed for a specific data set-up but it is also possible to apply the method and the approach to other data-sets using the same ratiocination.

In each of the steps in the pipeline various parameters affect the output and identified putative resistance genes, which parameters to use and tweak depends on the data and the aim of the analysis. The first thing is therefore to select the input data carefully. Data with distantly related species might not give meaningful homologous protein families using our cut-offs since proteins with similar function might be more different than our cut-offs allow. Step 1 in the pipeline combines the input data and creates a tables with homology count of all the cluster genes, in this step no filtering or selection is done and hence there is no parameters to tweak.

In step 2 selecting clusters with a specific homology count pattern, the strict or alternative patterns can be selected. Using the strict selection criteria, only 8% of the clusters meets the criteria. When using too restrictive selection patterns, there is a risk of creating false negatives, thus filtering away good cases. As mentioned earlier, many common secondary metabolite genes are likely to have homologs and belong to large protein families. Of the cluster genes belonging to this test dataset 12 and 6% belong to protein families larger than 102 and 153 proteins respectively, and these are found in 51% and 37% of the total clusters. Therefore using the ‘strict’ selection pattern, up to half of the clusters are likely to be discarded due to the homology count pattern of standard cluster genes, which causes a lot of false negatives. We therefore recommend using the alternative selection pattern. If the species in the dataset are distantly related the size of the protein families might be smaller and therefore a lower *Xinput* is recommended. The lowest reasonable value of *Xinput* we recommend is 1.5; going lower the risk of disregarding true resistance gene families becomes too big.

Filtering based on more clusters having the putative resistance gene (step 3) was designed to avoid false positives due to cluster prediction errors. This step is useful for data including closely related species likely to have similar clusters. If the data consists mainly of distantly related species, it is less likely that similar clusters will be found in more species and hence the risk of filtering away good cases increases thereby making false negatives. In the test data, about two-thirds of the cases are filtered away in this step, if running the pipeline and even more than two-tirds are filtered away, we suggest skipping this step.

The 4th filtering step was made to avoid false positives, in this step the percentage of organisms that should have a homolog of the putative resistance gene is selected and this parameter can be tweaked depending on the data. If the species are distantly related, it is more likely that some species have a different essential household mechanism and hence does not have a copy the household / target gene. A lower percentage of organisms that should have a homolog is thus recommended in this case. Another reason for choosing a lower percentage of organisms with a homolog is if the quality of the genome sequence data is low, with incomplete genomes or if the data includes novel species distantly related to model organisms where the gene prediction algorithms might not work as efficiently.

The last filtering step was also made to avoid false positives, in this step at least 50% of the species should only have a single homolog. This filtering step is independent of the quality of the data and the relatedness of the species, and we therefore recommend using this at all times. Here 30-60% of the cases are left after this filtering, significantly decreasing the number but leaving highly promising resistance cases.

## Conclusions

In this study, we have created a method identifying clusters responsible for producing bioactive compounds. The approach we have developed is based on a specific resistance mechanism and paves the way for rationally selecting promising bioactive clusters from whole genome data. The FRIGG pipeline was designed in connection with the *Aspergillus* sequencing project however several filtering steps and paramters can be tweaked to fit different kinds of data and to deal with the most likely errors from predictions and draft genomes.

We have tested the developed pipeline on 51 *Aspergillus* and *Penicillium* genomes identifying 72 unique putative resistance genes and clusters in the most strict configuration of the pipeline. In addition, the characterized Fellutamide B resistance gene *inpE* was successfully identified with this pipeline confirming the accuracy and applicability of the pipeline to such cases of resistance mechanisms.

As more and more genomes are sequenced, the relevance of this approach will increase and it will become a useful method for selecting which clusters to focus on in the hunt for novel drugs such as anti-fungicides, anti-cancer drugs and anti-microbial compounds.

## Materials And Methods

### Fungal species

The data consisted of 50 *Aspergillus* and 1 *Penicillium* species with available whole genome sequencing data which was downloaded from JGI. Species information can be found in Additional Table 5.

### Input data

The data used in the pipeline was organized in a MySQL database, an overview of the input data can be found in the file: Input_data_pipeline.txt and the data can be found as sql/csv files on https://github.com/ingek-1/FRIGG_pipeline. A few of the data files were too big for Github and can be found in this repository instead https://files.dtu.dk/u/ox6SPjaekiyxHpI8/Data_tables_FRIGG?l. The data includes predicted secondary metabolite gene clusters based on an implementation of the SMURF (21) pipeline described elsewhere (6). Homologous protein families were created with Aspergillus optimized parameters based on single linkage of bidirectional BLASTp hits with identity ≥ 50% and sum of query and hit coverage ≥ 130% as described in (6). InterPro annotations of the proteins were also used (19, 20) and gff information.

### Pipeline

The pipeline was created using Python and the investigation of specific protein families was conducted in R(28). The versions and packages used can be seen in version info files on the github page. For alignment and trimming of the protein families clustalo and Gblocks was used in a Python script, also included on the github page. All the scripts and files are available at Github: https://github.com/ingek-1/FRIGG_pipeline.

## Acknowledgments

M.R.A. and T.V. gratefully acknowledge funding from the Villum Foundation, Grant VKR023437. The funders had no role in study design, data collection and interpretation, or the decision to submit the work for publication.

Uffe H. Mortensen and Jens C. Frisvad are gratefully acknowledged for their input on the initial hypothesis and requirements of the pipeline.

## Supplemental Material

Guidelines for supplemental material appear in the Instructions to Authors. This section of the paper should include legends for any supplemental material that is intended for posting. Such supplemental material must be submitted with the manuscript. Files can be added to the submission at the publisher’s submission site. Here is a list of sample legends for supplemental material:

**FIG S1 Common InterPro domains in secondary metabolism**. Visualization of the most common InterPro annotations of secondary metabolite genes (found in more than 1000 secondary metabolite proteins) and the size of the protein families. Two horizontal lines indicate the recommended protein family size cut-offs where *X_Input_* is 2 (102) and 3(153).

**FIG S2 Principal component analysis of the protein family 597268**. Principal component analysis of the protein family 597268 containing two potential resistance genes (protein id 11595 and 32200) found in *A. oryzae* and *A. flavus*. The panels to the left are colored based on the sections the proteins belong to while the panels to the right are colored based on if the protein if found in a a selected cluster (StrictClust-green), not in a cluster (0-blue), and the homolg to the ones found in selected clusters (outsideSC-yellow).

**FIG S3 Phylogenetic tree of the the protein family (597268)**. Phylogenetic tree of protein family 597268 containing two potential resistance genes (protein id 11595 and 32200) found in *A. oryzae* and *A. flavus*. The node labels shows bootstraps values based on 500. The tip labels have the species name, the section, protein id, number of homologs in the species and an indication if it is found in a selected cluster (StrictClust), a predicted cluster (Clust), not in a cluster (0) and the homolg to the ones found in selected clusters (outsideSC)

**TABLE S1**. Species used in this study, showing species name, section, and link to the JGI pages with the genomes.

**TABLE S2**. Overview of the 72 identified putative resistance genes families and the parameters where they were identified.

